# Active and Allosteric Site Binding MOLECULAR MECHANICS-QUANTUM MECHANICS studies of STEVIOSIDE Derivative in PCSK9 Protein Intended to provide a safe Antilipidemic agent

**DOI:** 10.1101/2023.05.04.539221

**Authors:** N Irfan, Prakash Vaithyanathan, Harishchander Anandaram, S Mohammed Zaidh, S Priya Varshini, A Puratchikody

**Author notes:** <u>Corresponding Author:</u>* **Prakash Vaithyanathan**, *E-mail:.

## Abstract

Interaction of low-density lipoprotein receptors with proprotein convertase subtilisin/ kexin type 9 (PCSK9) plays a vital role in causing atherosclerosis. It is the hidden precursor of clinical myocardial infarction (MI), stroke, CVD and estimates 60% of deaths worldwide. The current need is to design small molecules to prevent the interaction between PCSK9 with LDL receptors. This study aims to evaluate the PCSK9 antagonistic effect of a derivative of Stevioside (also referred as Methylidene tetracyclo derivative) and atorvastatin. Also, a comparative study was performed to analyze the binding interaction of molecules inside the active and allosteric sites of PCSK9. The RCSB downloaded protein 7S5H and above said ligands were optimized to the local minima energy level and docked inside the active and allosteric sites. The stability of non-bonded interaction of complex was analyzed using Desmond MD simulation studies. The results of docking showed that the Methylidene tetracyclo molecule possesses a two-fold higher affinity of -10.159 kcal/mol in the active site and -10.824 kcal/mol in the allosteric site. The Phe377 amino acid made the Methylidene tetracyclo molecule orient inside the active site. Nine H-bonds with 5 amino acids of allosteric site increase the binding affinity compared to Atorvastatin. The MD simulation studies exposed that the nonbonded interaction of Methylidene tetracyclo molecule was stable throughout 100ns. This confirms the Methylidene tetracyclo molecule will be the better hit as well as the lead molecule to inhibit PCSK9 protein.

## 1. Introduction

In the blood, cholesterol is unique and key precursor for triglycerides, phospholipids, and all steroids (Bhattarai et al., 2021) as well as participating in catabolism of important hormones such as progesterone, estrogen, adrenal corticosteroids and testosterone. This vital molecule becomes harmful while crossing the limits in blood (hyperlipidemia) and these lipid particles stick to the vessel of blood flow and blocks the run, which causes risk of stroke and heart attack. This condition of very high levels is named as atherosclerosis (Russo & Jang, 2022; Seidman et al., 2014) Anti-cholesterol drugs are used in this condition whenever the person suffers from hyperlipidemia.

Usually, statins are the drugs used for this condition for 30 years. The first statin lovastatin (Mevacor) was permitted for commercial use in the USA. In the late 1980s, statins reduce the risk of further heart attacks in people who’d already had one. (Basak & Basak, 2020) Some of the benefits to take statins include cholesterol reduction, cardiovascular function, and inflammation reduction. However, the side effects of these drugs are considered the primary reasons that people may stop taking statins. (Ahmad, 2014) One of the drugs cerivastatin withdrawn from the marketplace for the reason that cause 52 deaths. Also, it linked to rhabdomyolysis that led to failure of renal system. (Furberg & Pitt, 2001) Alternatively, Ezetimibe is a ‘inhibitor of cholesterol absorption’ that confines the captivation of cholesterol from GIT region. (Lloyd-Jones et al., 2022) Market size of statin was grasped US$ 14.9 Billion in 2022 globally with CAGR growth rate of 3.2% during 2023-2028.

The catalytic domain of PCSK9, which is similar to subtilisin, can bind to the LDLR EGF-A (epidermal growth factor-like repeats) domain. Asp-374, which is known to be altered to Tyr and increases PCSK9’s (McDonagh et al., 2016; *PCSK9 Inhibitors - PubMed*, n.d.; Schmidt et al., 2020; Zainab et al., 2021)affinity for LDLR, is found in this domain. In 2021. To reduce cholesterol buildup and the risk of coronary artery disease, the binding of LDL to LDLR is crucial. The association of PCSK9 with LDLR enhances cholesterol buildup(Du et al., 2011) by preventing LDL from attaching to LDLR. In order to avoid CAD, it is crucial to suppress PCSK9 and its interaction with LDLR. Small-molecule (Kamiya et al., 1979)pharmacotherapies and biological therapies using monoclonal antibodies are the two primary approaches used in PCSK9-targeted treatments.

Since the PCSK9 molecular structure necessary for small molecule binding to decrease this protein activity(Zainab et al., 2021) has not been localized, progress in the discovery of small-molecule pharmacotherapy for PCSK9 inhibition (Xu et al., 2019)is unfortunately challenging. In 2019b, Xu et al. Computing methods based on bioinformatics and cheminformatic(Galvão Lopes et al., 2023; Malla et al., 2022; Rehman et al., 2022; Sattarinezhad et al., 2016)s may be useful for locating novel binding sites for the inhibition of a protein. Cheminformatics and bioinformatics deal with developing and modifying databases and statistical methods to manage and analyzed biological and chemical data. Their equipment may be used to locate and examine active sites, macromolecular biological structures, and drug targets.(Irfan et al., 2022) Using a Computational Fragment-Based Method for Protein Binding Site Analysis for Drug Discovery In this work docking and dynamic simulation techniques used to study the binding profile of the Methylidyne tetracyclo derivative and realistic model-based validation also performed to analyses the behavior of the molecules in Active site and allosteric site.

## 2. Materials and methods

### 2.1 Software and Modules

To study the atomic level interactions and stability of the complex between ligands and proteins, Schrodinger, BIOVIA Discovery studio, and Desmond simulation software were used. The modules protein preparation, Grid generation, Site map prediction, Ligprep, and glide docking modules were used to analyze the molecular mechanic’s interaction of Methylidene tetracyclo Derivative with PCSK9 (Abboud et al., 2007; Du et al., 2011; Herbert et al., 2010; Xu et al., 2019)protein, and dynamic simulation, simulation quality, simulation interaction diagram modules were used to study the QM-based stability studies of the complex(Bitencourt-Ferreira et al., 2019; Kuki & Nielsen, 2010).

### 2.2 Protein and ligand preparation protocol

To study the interaction between the 7S5H protein, Atrovostatin, and methylidene tetracyclo has to be prepared as a stable molecule with a local energy level. The protein 7s5h was given as source entry in the preparation protocol and the preprocessing of protein was performed with the setup of Assign bond order using the CCD database and the hydrogen are replaced. The metal, disulfide bonds are created based on the zero-order. The Epik generates het states pH set as N.N ±2.0. this preprocessed protein was subjected to the optimized H-Bond assignment with the setting of PROPKA optimization technique with sample water orientation. Finally, the protein was minimized with heavy atom RMSD coverage of 0.30 A and the OPLS4 force filed calculation. The water presents away from the PDB ligand in the distance of 5 A was deleted. The methylidene was tetracyclo and Atrovastin molecules were prepared for docking using the LigPrep protocol. The algorithm used the OPLS4 force field parameters with Epik and the system was desalted. Tautomers and 32 stereoisomers are computed per ligands. (Madhavi Sastry et al., 2013)

### 2.3 GRID generation and binding pocket analysis

The pdb protein 7S5H possesses the peptide molecules and the receptor grid generation protocol considered the site as a binding pocket of the methylidene tetracyclo and Atrovastin. Van der Waals radius scaling factor is set as 1.0 with a partial charge cutoff of 0.25. The enclosing box was fit in the centroid of workspace ligand and length of dock ligands was fixed to 12 A. Site Map protocol was used to detect the Allosteric site in the 7S5H, standard grid with 4A crop site maps at the nearest site point. Based on the site map points allosteric grid is generated to dock the molecules.(Wang et al., 2013)

### 2.4 Ligand docking

The ligand docking protocol was set up with a Scaling factor of 0.80 and a partial charge cutoff of 0.15 for van der Waals radii. The grid file was loaded and the XP extra precision setup with the flexible ligand sampling. Sample nitrogen inversion, rig conformations considered, and all predefined functional groups used to bias sampling of torsions. Epik state penalties were added to the final dock score. The energy window for ring sampling is fixed at 2.5 kcal/mol for conformer generation and the minimization distance-dependent dielectric constant value is 2.0. (Malla et al., 2022)

### 2.5 System Builder and Dynamic simulation set up

The best-docked pose of Methylidene tetracyclo(Pincock & Torupka, 1969) Derivative and Atrovastatine stability was studied using the above-titled protocol. Initially, the complex was solvated using the pre-defined SPC solvent model, and the boundary condition was set up as orthorhombic box shape, buffer box size calculation method used in the distance of 10.0 A. Also, the final volume is minimized based on the surface occupation of complex protein and the system was neutralized by adding Na+ and salt negative ions (Cl-). The solvated model was submitted to molecular dynamic simulation protocol. Simulation time was fixed to 100 ns and trajectory recording interval of 100 ps with a 1.2 energy gap. Totally 1000 frames are generated in an NPT ensemble class temperature of 300 k with 1.01325 bar pressure and the model was relaxed before simulation.

### 2.6 System quality analysis and dynamic interaction diagram

The final complex simulation with loaded trajectory quality was analyzed block length for averaging 10.0 ps. This simulation quality analysis protocol summarizes the 100ns simulation and plots the graphs using the Total energy (kcal/mol), Potential energy (kcal/mol), Temperature (k), Pressure (bar), and volume (A3) parameters. Eventually, the simulation interaction diagram module calculated the RMSD, RMSF, and Torsion values of a complex in the 100ns time scale.

## 3. Result and Discussion

### 3.1 Optimization and Grid Box

The protein 7S5H and ligand molecules preparation protocols are optimized to the local minima to make it stable for the docking studies. Figure 1 illustrates the superimposed image of protein before and after minimization. The major conformational changes observed near the binding region loop amino acids of ASP192, ASP238, and GLY 384. Similarly, the fragment of the molecule Methylidene tetracyclo derivative (Pincock & Torupka, 1969)superimposed with the before minimized molecules. The configurational changes were observed in the fragments of the cyclic ring with the RMSD of 0.4570. the final energy found to be the molecules was 86.852 kcal/mol and the protein was -2272.338 kcal/mol.

**Figure 1.**
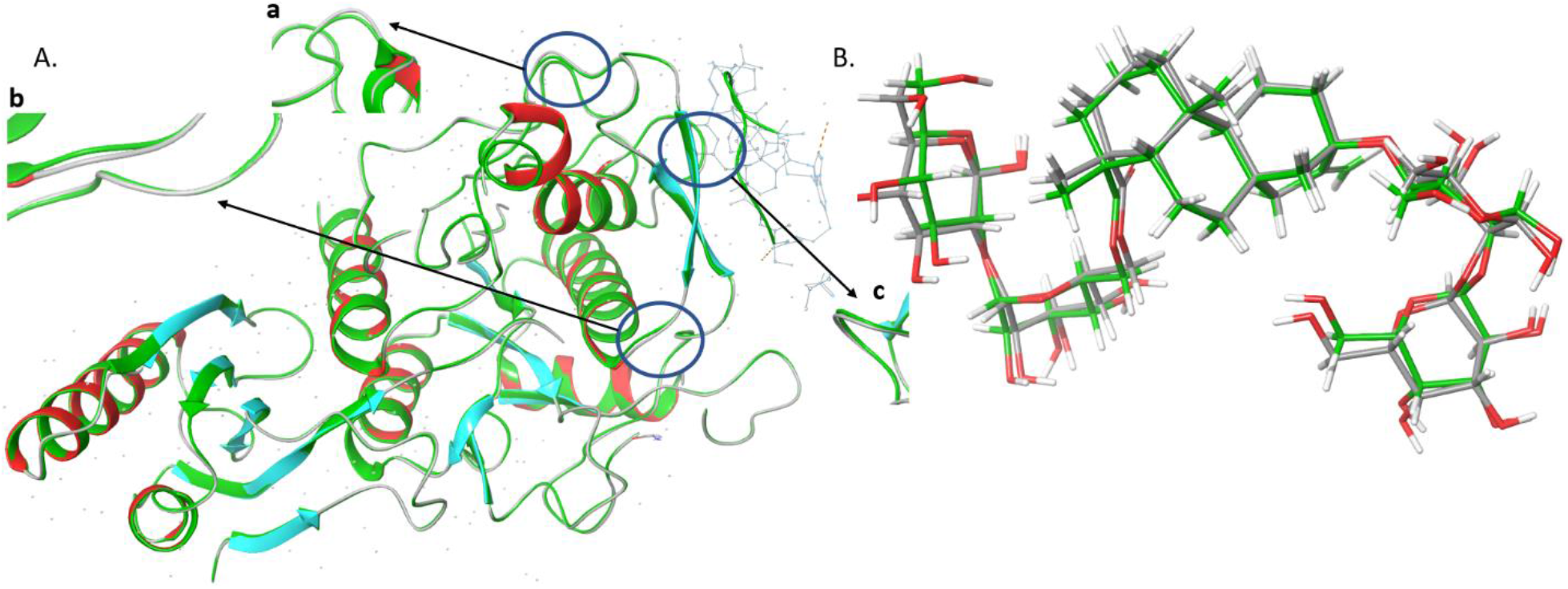
**A**. Superimposed Secondary structure of protein 7s5h before (green) and after (red) minimization. B. Superimposed structure of Methylidene tetracyclo derivative before (green) and after (red) minimization.

Two binding pockets such as the active site and the allosteric site were selected to generate the grid to bind the molecules. Figure 2A illustrates the grid box position in the 7S5H protein With the coordinates of X -17.16, Y 19.08, and Z -6.2. Similarly, the binding site detection protocol predicts the best allosteric site in the coordinates of x -22.5, Y -20.67, and Z 11.3 and a radius of 10 A.

**Figure 2.**
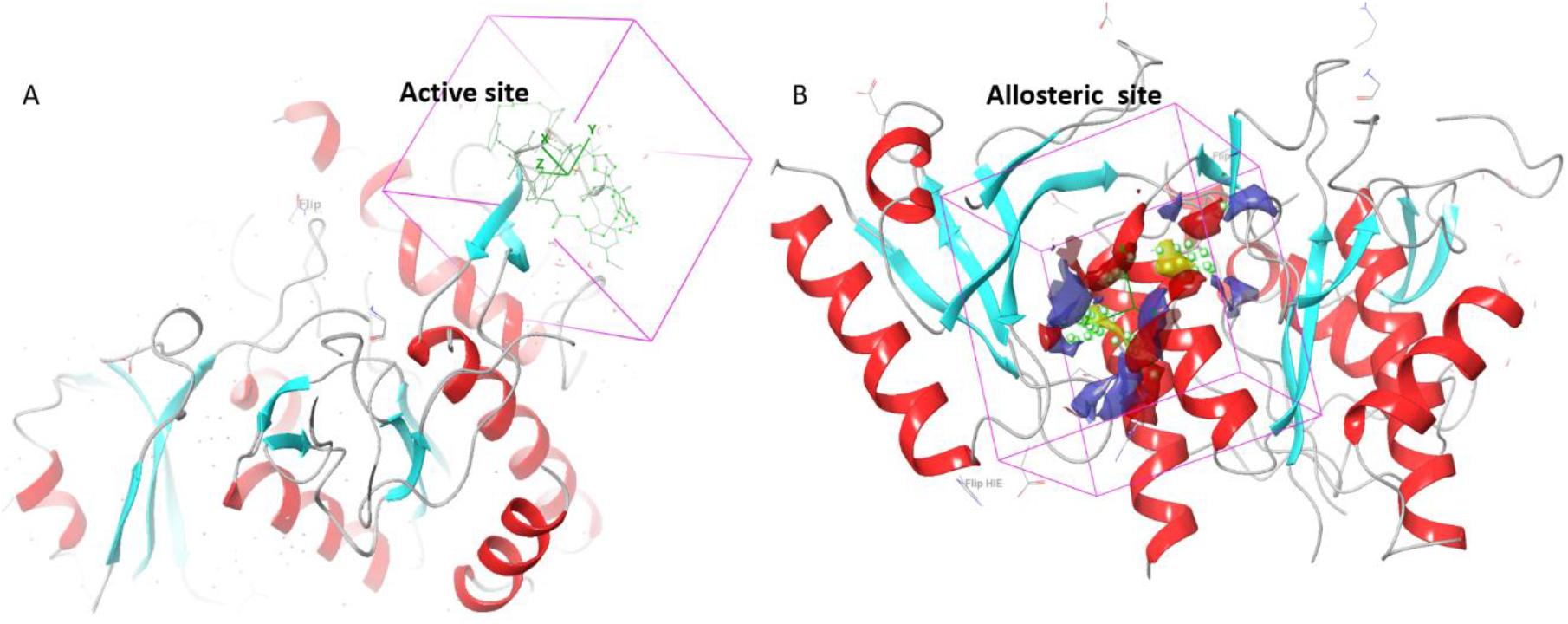
**A**. Active site of 7s5h protein. B. Allosteric site of 7s5h protein.

**Figure 3.**
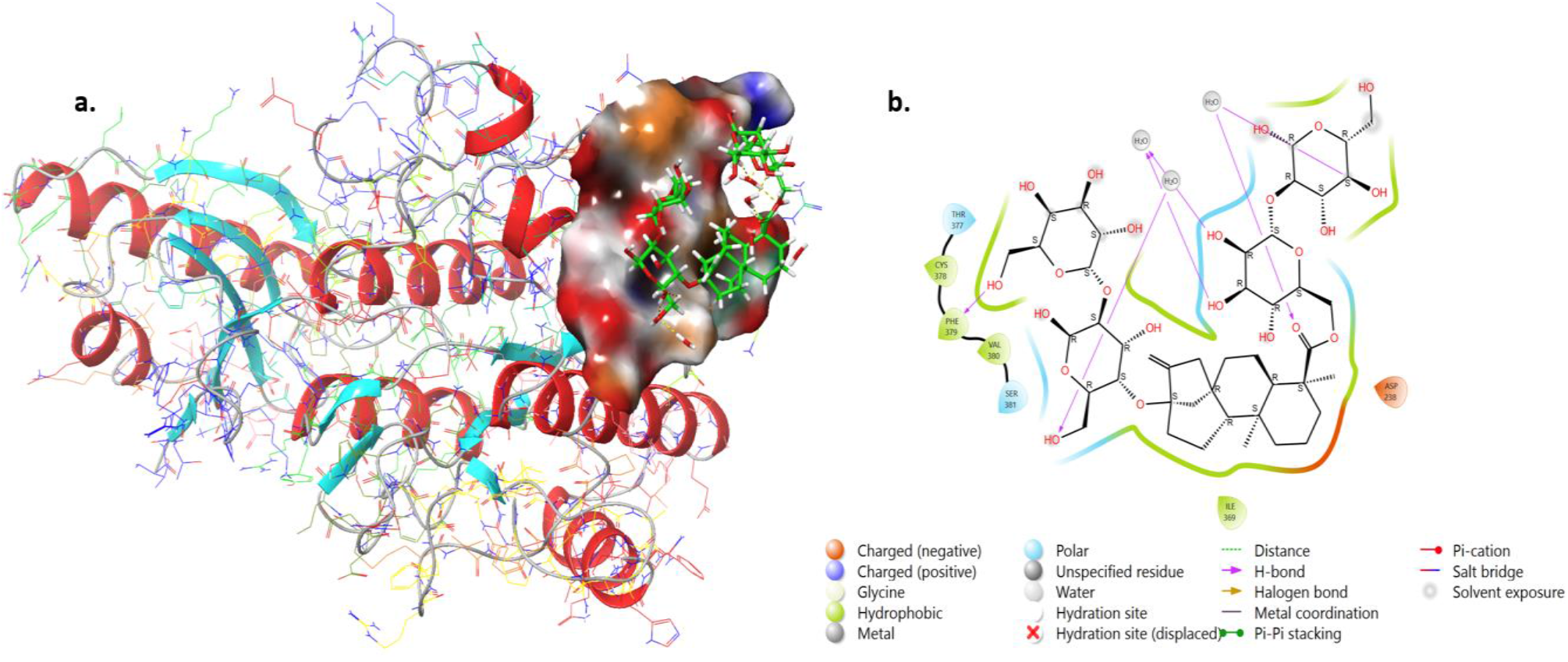
**a**. Binding of Methylidene tetracyclo derivative in the PDB active site of 7s5h protein. b. 2D interacting amino acids with the atoms of Methylidene tetracyclo derivative.

**Figure 4.**
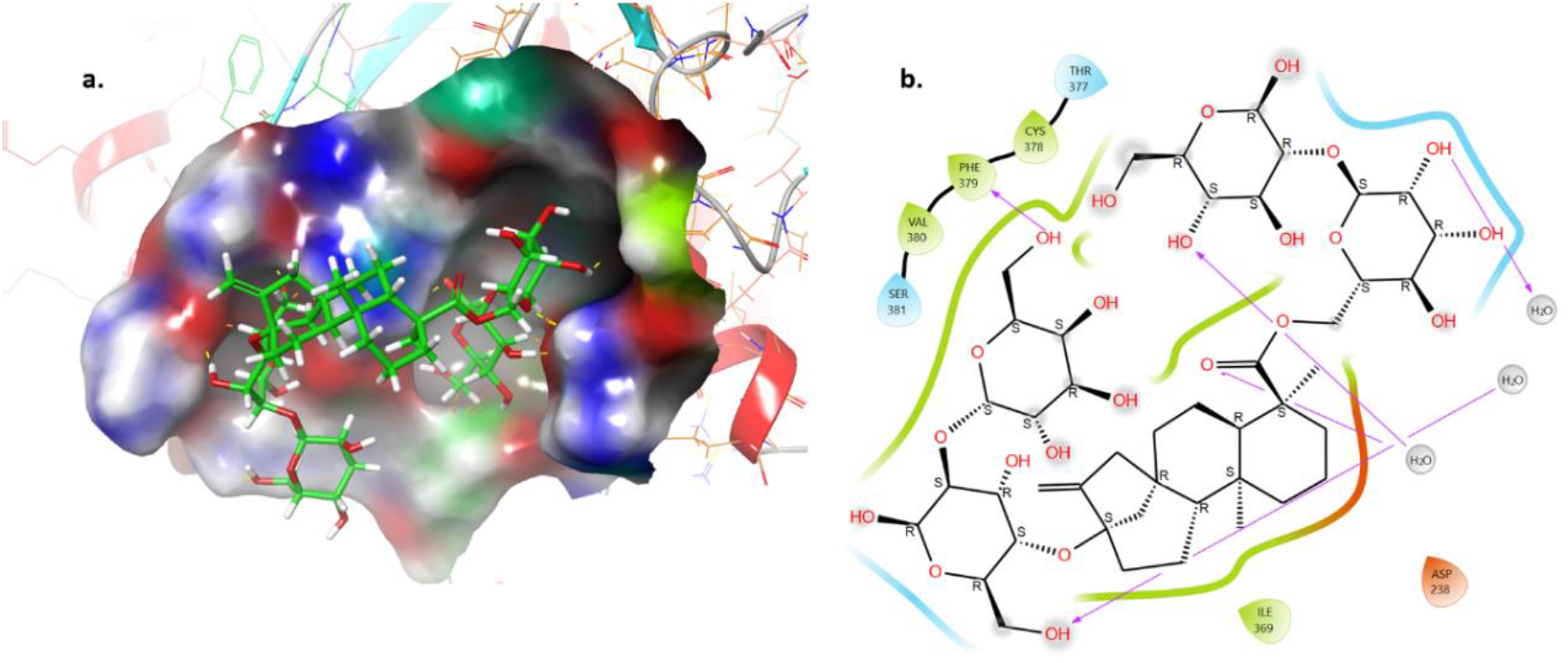
**a**. Binding of Methylidene tetracyclo derivative in the Allosteric site of 7s5h protein. b. 2D interacting amino acids with the atoms of Methylidene tetracyclo derivative.

### 3.2 Ligand docking

The results of glide docking showed that, both the molecules Methylidene tetracyclo derivative and Atorvastatin formed good possess inside the binding pocket. The energy of the Methylidene molecule was two folds higher in both sites -10.159 kcal/mol and -10.824 kcal/mol respectively. The water molecules aiding methylidene molecule binding better compare to the standard and -OH groups of the molecule generated 5 H-Bonds with a water molecule and one H-bond with Phe379 amino acid and pyran substituted OH group. Additionally, 6 amino acids Ser381 and Thr377 (polar), Val380, Cys 378, and Ile 369 (Hydrophobic and Asp238 (Negative charge) change the configuration of the Methylidene derivative to bind well.

The allosteric binding pocket (Ludington, 2015)also produced multiple links without the support of the water molecules. Nine H-bond interactions linked with ThrA:63, ThrA:61, AspA141, ArgB295 AsnB:298, and GlnB 302. Which coiled the molecule to make proper confirmation to interact with other amino acids through the different interactions such as Hydrophobic (AlaA62, TyrA142, ValA81, LeuA119, ProB331, AlaB330, AlaB299, TyrB293), Positive charged (ArgB295, Lys A83, Arg B306), Polar (ThrA61, ThrA63, HisA65, GlnB302, AsnB298) and Negative charged (AspA141, GluA84, GluB332).

A two-dimensional interaction analysis of the standard drug atorvastatin inside the active site and the allosteric site was shown in Figure 5. The image exposed that standard atorvastatin formed four types of interaction and the carboxylic group and aliphatic chain OH group were involved in the hydrogen bond interaction. An amino acid Ser381 interacts with the R-configured OH group bond and the Asp238 negatively charged interaction supports the aniline ring orientation. The amino acid Phe379 linked with benzene ring system of the molecule by π-π interactions helps the pyrrole ring position in the active site (Figure 5a).

**Figure 5.**
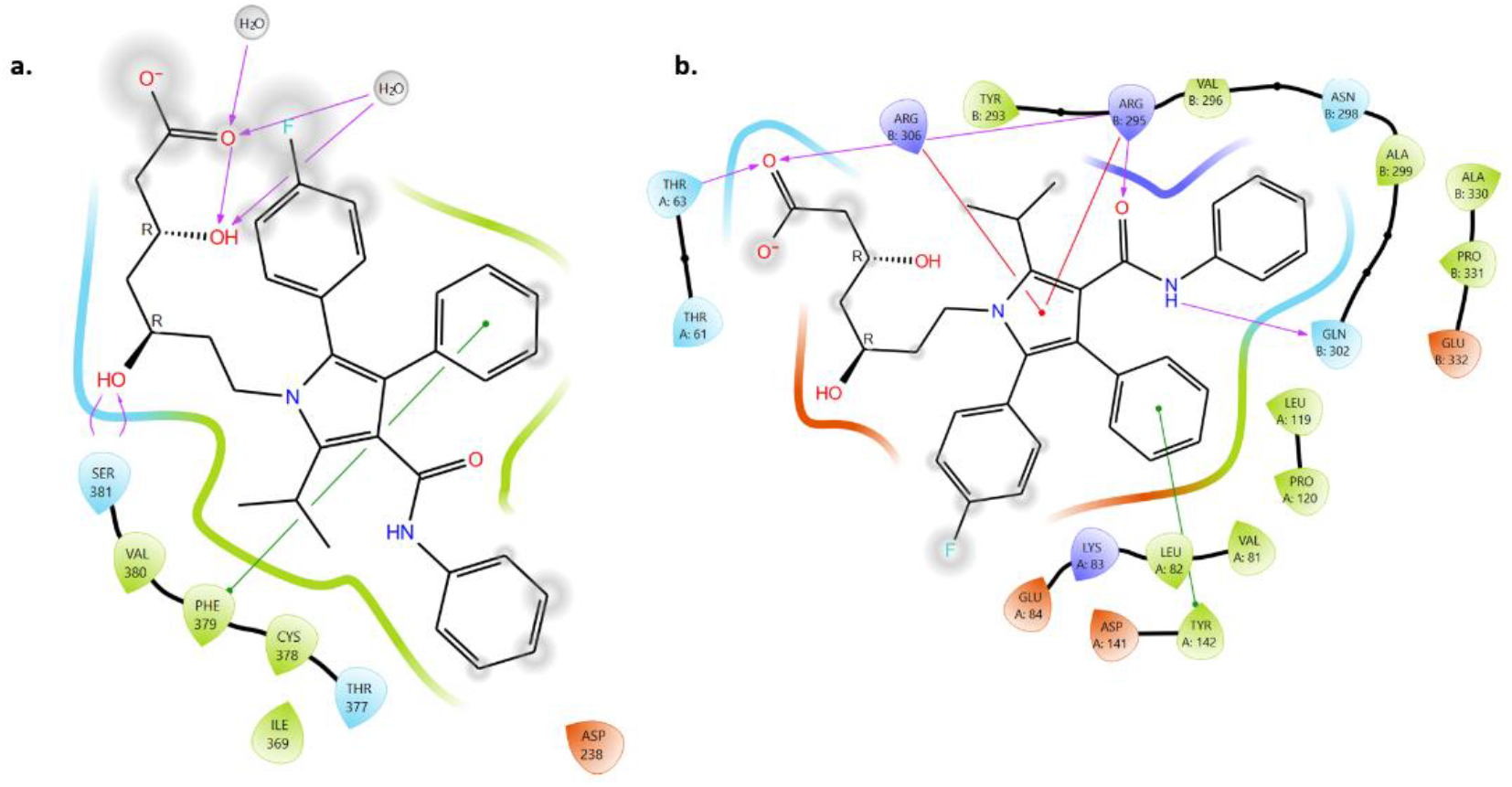
**a**. Binding of Atrovastatine in the active site of 7s5h protein. b. Binding of Atrovastatine in the allosteric site of 7s5h protein

**Figure 6.**
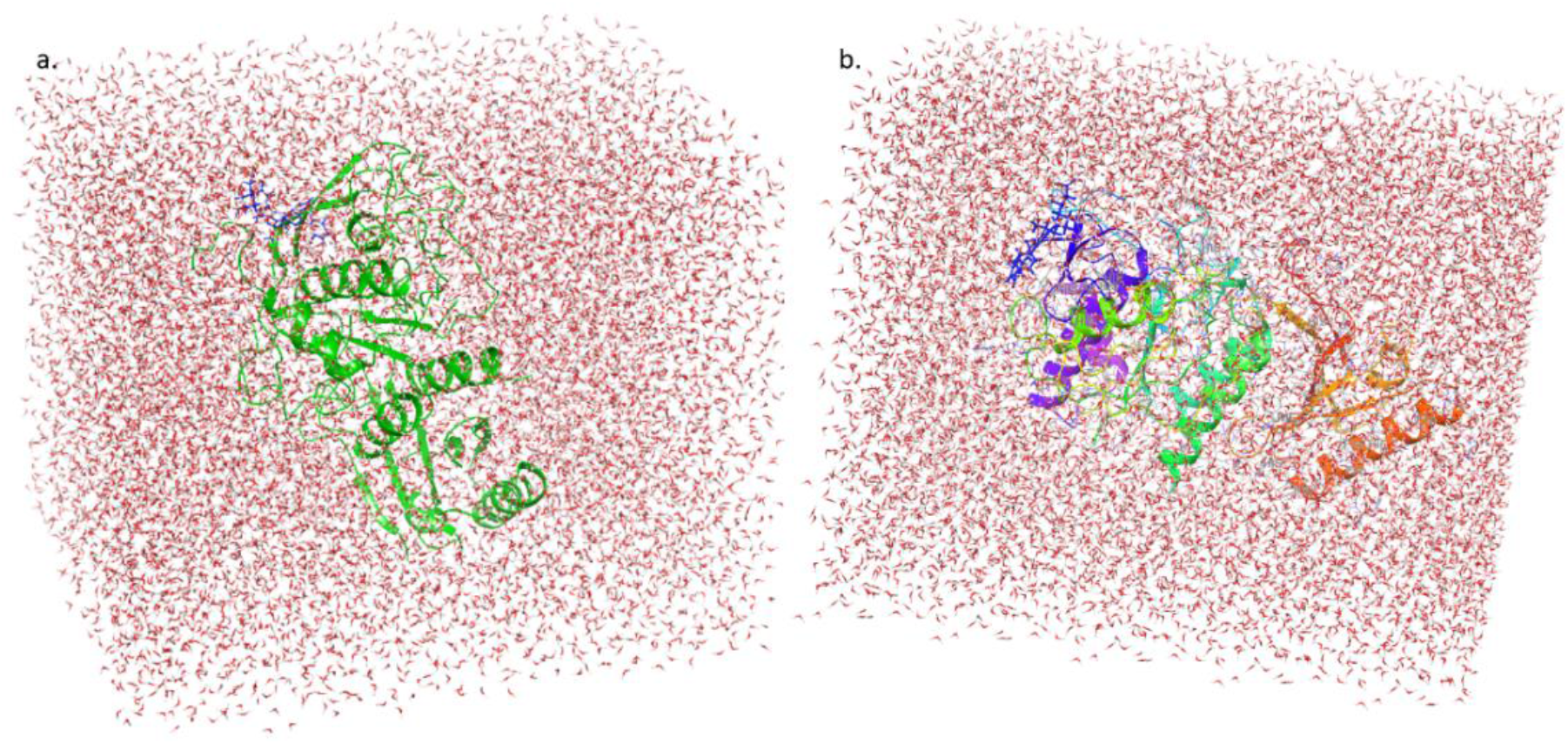
**a**. Solvated model of Active site binding of Methylidene tetracyclo derivative. b. Solvated model of Active site binding of Atorvastatin

The allosteric binding of atorvastatin is shown in figure5b, which exposed that H-bond and π-π interactions significantly played the role in allosteric binding. Hydrophobic amino acids AlaB299, and polar amino acids AsnB298, and Gln302 support Aniline ring fixing, LeuA82, ValA81, ProA120 and LeuA119 aid the benzene ring position, and ArgB295, ArgB306 formed π-cationic fixes the core center pyrrole moiety orientation inside the allosteric site (Figure 5b).

### 3.3 Dynamic simulation stability analysis

Interaction analysis alone is not enough to confirm the possible effectiveness and bond formation by the Methylidene tetracyclo derivative. It should be analyzed for dynamic stability using MD simulation studies. (Galvão Lopes et al., 2023; Navabshan et al., 2021; Petrilli et al., 2020)The Methylidene tetracyclo derivative and Atorvastatin molecule complexes were solvated with water.

The system builder protocol added 12355 water molecules to the active site binding complex of Methylidene tetracyclo and 12350 water molecules for the Atorvastatin protein complex system for 100ns simulation studies.

### 3.4 System quality analysis and dynamic interaction diagram

The molecular dynamic simulation run quality was checked and the parameter values be good in range (Table 2). Figure 7 illustrates they were found to be good and the complex simulation quality has stable volume, pressure, Temperature, Potential energy, and energy (Figure 7)

**Table 1.** Dock score of molecules in the active site and allosteric site of the 7s5H protein

| Name of the molecule | Docking Energy of Active site (kcal/mol) | Docking Energy of Allosteric site(kcal/mol) |
| --- | --- | --- |
| Methylidene tetracyclo derivative | -10.159 | -10.824 |
| Atrovastatine | -3.953 | -4.607 |

**Table 2.** Simulation Quality of Active site run of Methylidene tetracyclo protein complex run

| **Quality values of Methylidene tetracyclo protein complex simulation (100ns)** |  |  |  |
| --- | --- | --- | --- |
| **Properties** | **Average** | **Std.Dev** | **Slope (ps<sup>-1</sup>)** |
| Total energy (kcal/mol) | -110704.044 | 100.376 | -0.001 |
| Potential energy (kcal/mol) | -136457.605 | 87.838 | -0.001 |
| Temperature (K) | 298.704 | 0.576 | 0.000 |
| Pressure (bar) | 1.517 | 49.395 | -0.000 |
| Volume (A3) | 423742.763 | 396.905 | -0.001 |

**Figure 7.**
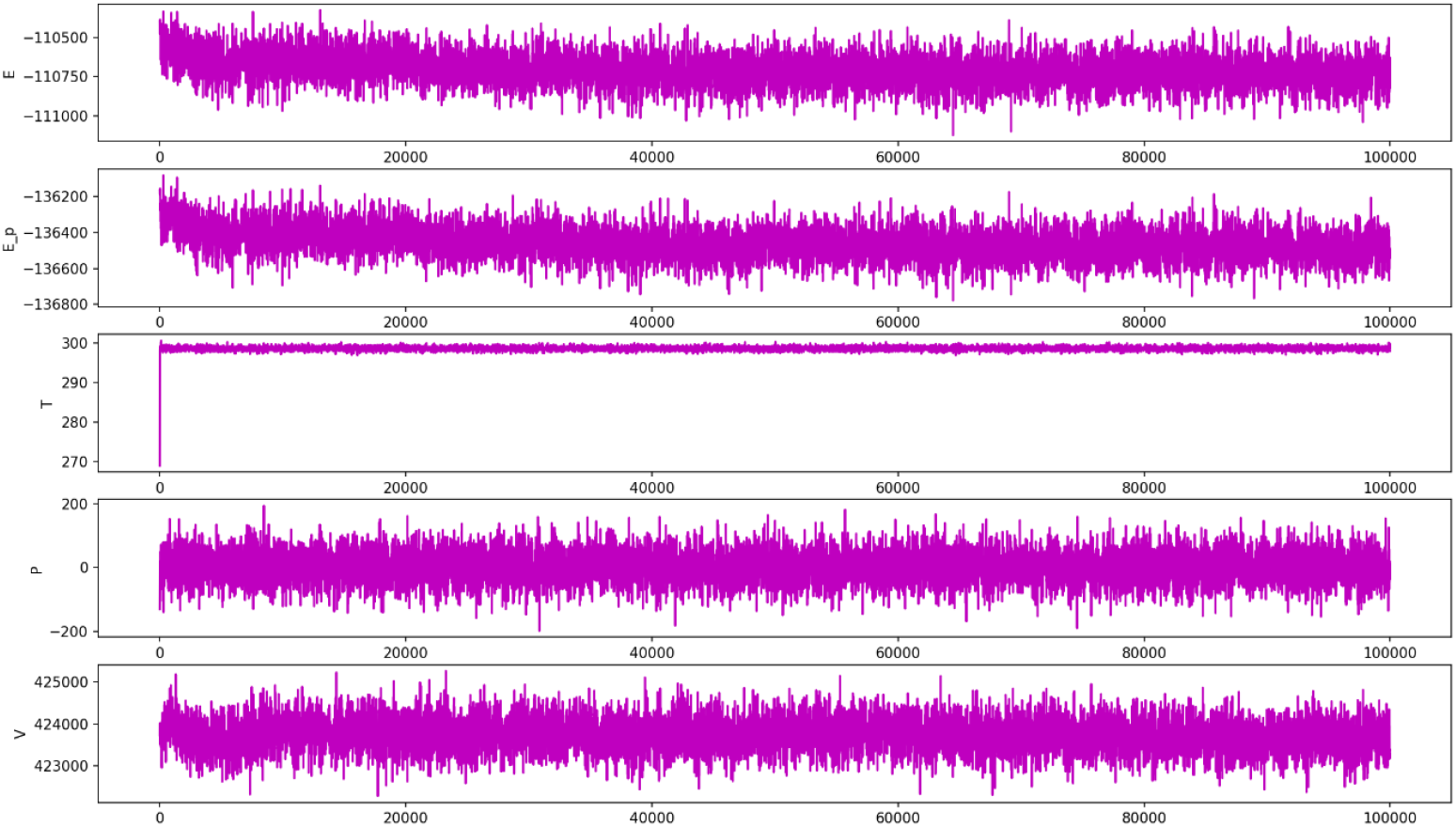
Quality analysis report of the 100 ns simulation run of Methylidene tetracyclo protein complex.

### 3.5 RMSD, RMSF and Protein ligand Contacts in active site

The RMSD graph showed that the 7s5h protein stats stabilize at the nano seconds of 32 with the Cα chain of complex (7S5H-Methylidene tetracyclo) RMSD range of 1.52 A – 1.85A Figure 8a. The Atorvastatin-7S5H complex RMSD was shown in figure 8b, which explain that the complex is not stable up to the 80ns.

**Figure 8.**
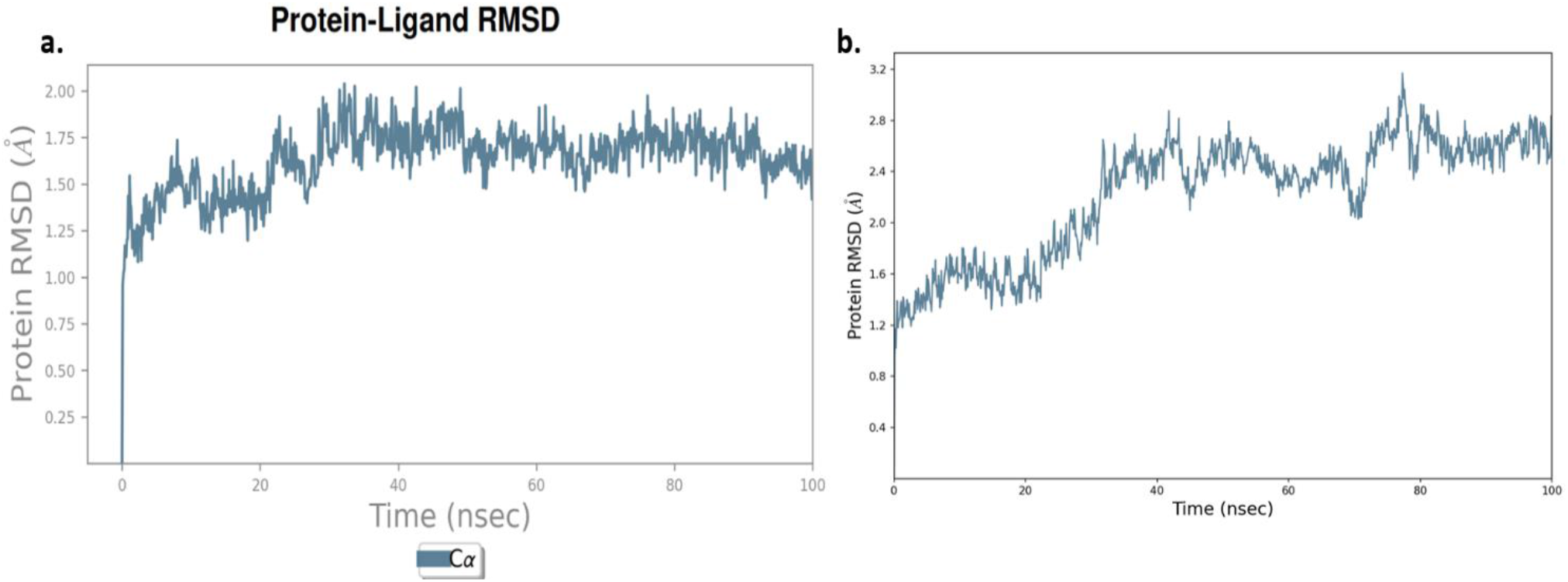
a. RMSD of different confirmation of 7S5H-Methylidene tetracyclo at 100ns. b. RMSD of different confirmation of 7S5H-Atrovastatin at 100ns.

The 7S5H-Methylidene tetracyclo complex contacts showed in the Figure 9, in the simulation run additional interaction formed and aide the molecule to completely stable up to the 100ns time course. Negative charged Glu 84, 85, 366,410 and Asp 367 interaction additionally generated the interaction during the simulation. The polar amino acids Thr86, Ser381, Thr 377, Gln 413 and Hydrophobic interaction Ala341, Ile369 and Phe379 orient the rigid ring inside the active binding pocket of 7s5h protein. These amino acids interactions and configuration reduce the fluctuations of complex and stablish it.

**Figure 9.**
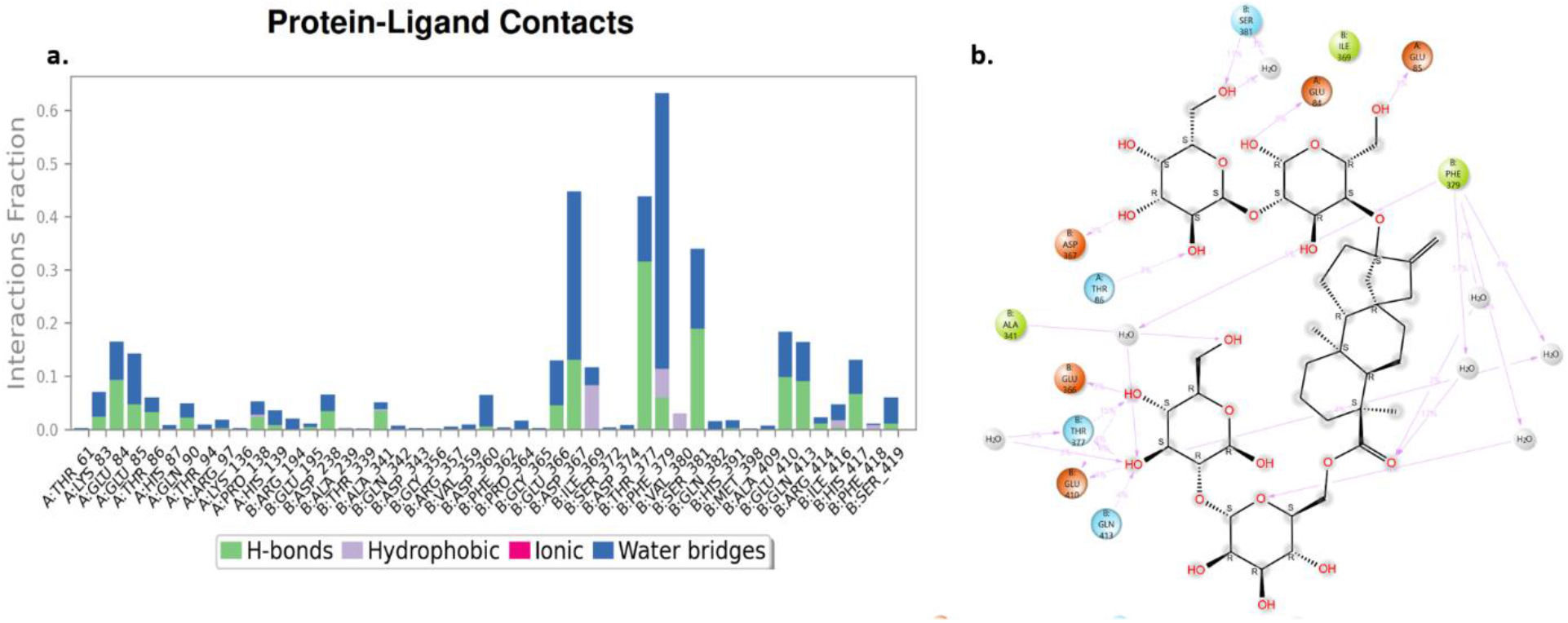
**a**. Methylidene tetracyclo derivative contacts during the 100ns simulation. **b**. Interaction map of the Methylidene tetracyclo molecule.

Figure 10 illustrates the higher RMS fluctuation of amino acids observed in the atorvastatin-protein complex compared to the RMSF of Methylidene tetracyclo derivative. Specifically, the amino acid 132, 143, 250 fluctuated more than 5 A and leads the less stability of atorvastatin with 7S5H protein.

**Figure 10.**
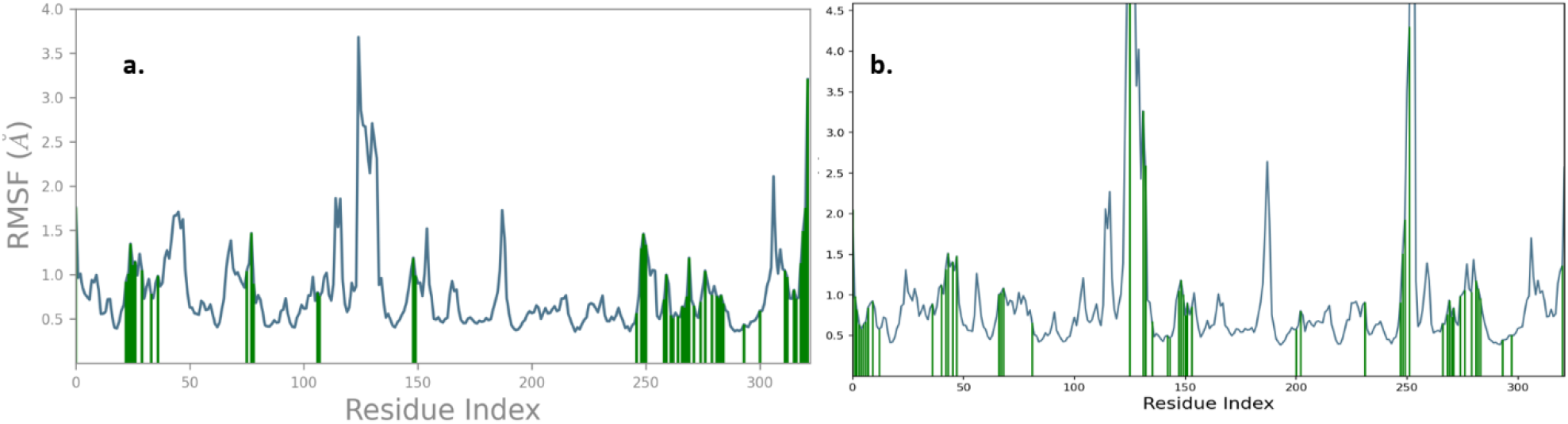
a. RMSF of Methylidene tetracyclo derivative-7S5H protein. b. RMSF of Atorvastatin.

### 3.6 RMSD, RMSF and Protein ligand Contacts in allosteric site

Stability of the Methylidene tetracyclo derivative and the atorvastatin was analyzed and illustrated in the following figures. RMSD(Alder & Wainwright, 1959; Doshi & Hamelberg, 2015; Paquet & Viktor, 2015; Shukla & Tripathi, 2020a, 2020b) of allosteric binding molecules explain the complex start getting stablish from 26ns and little fluctuated in the 46 ns and made permanent stability (Figure 11a) and the deviation range is 1.48-1.90 A. The Atorvastatin (Figure 11b) molecule not getting stabilize up to 71ns with deviation of 2.0-2.8 A.

**Figure 11.**
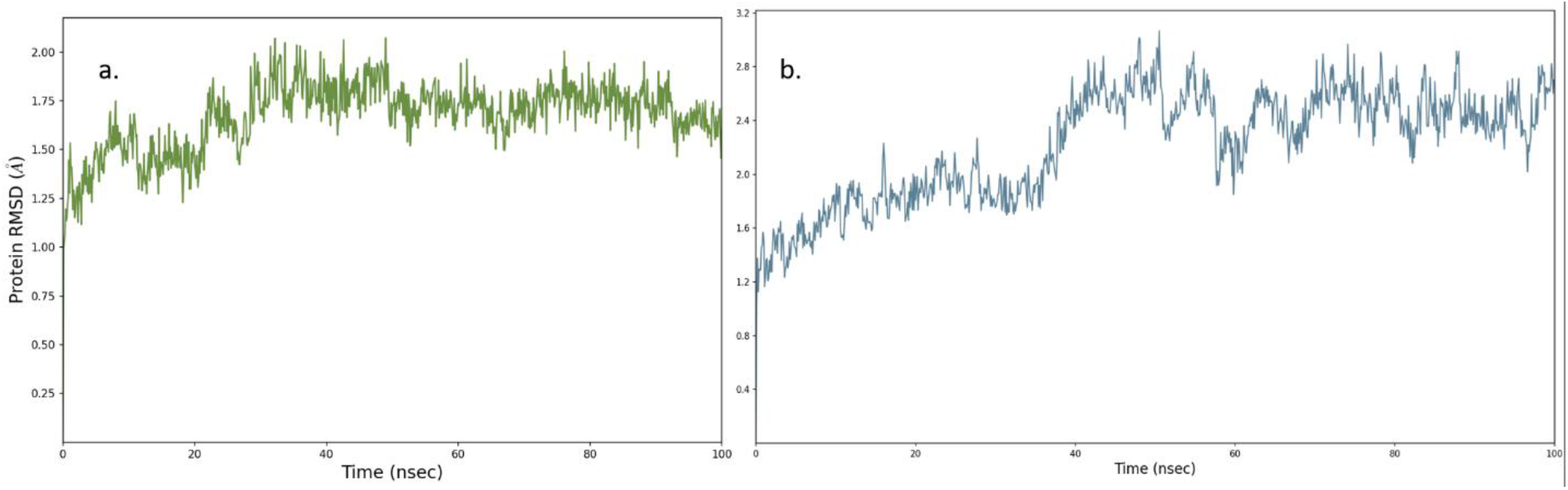
a. RMSD of different confirmation of 7S5H-Methylidene tetracyclo at 100ns inside the allosteric site. b. RMSD of different confirmation of 7S5H-Atrovastatin at 100ns inside the allosteric site.

Totally 32 residues involved in the 7S5H-Methylidene tetracyclo derivative allosteric binding during the simulation. Which reduces the confirmational changes of the side chain hydrogen to make stable confirmation of the complex (Figure 12a). Specifically, B chain residue Thr377, Ser381plays vital part in folding stabilization of protein. In the atorvastatin binding few amino acids only involved in the stabilization of protein (Figure 12b).

**Figure 12.**
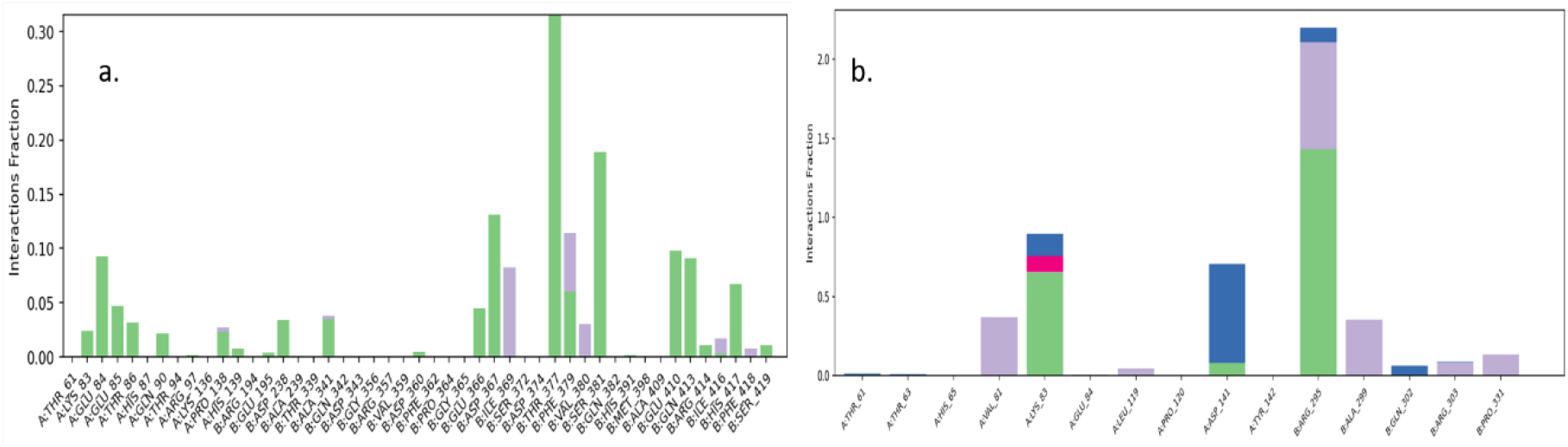
a. Allosteric site interaction formed during the 100ns simulation of 7S5H-Methylidene tetracyclo derivative. b. Allosteric site interaction formed during the 100ns simulation of 7S5H-Atrovastatin.

**Figure 13.**
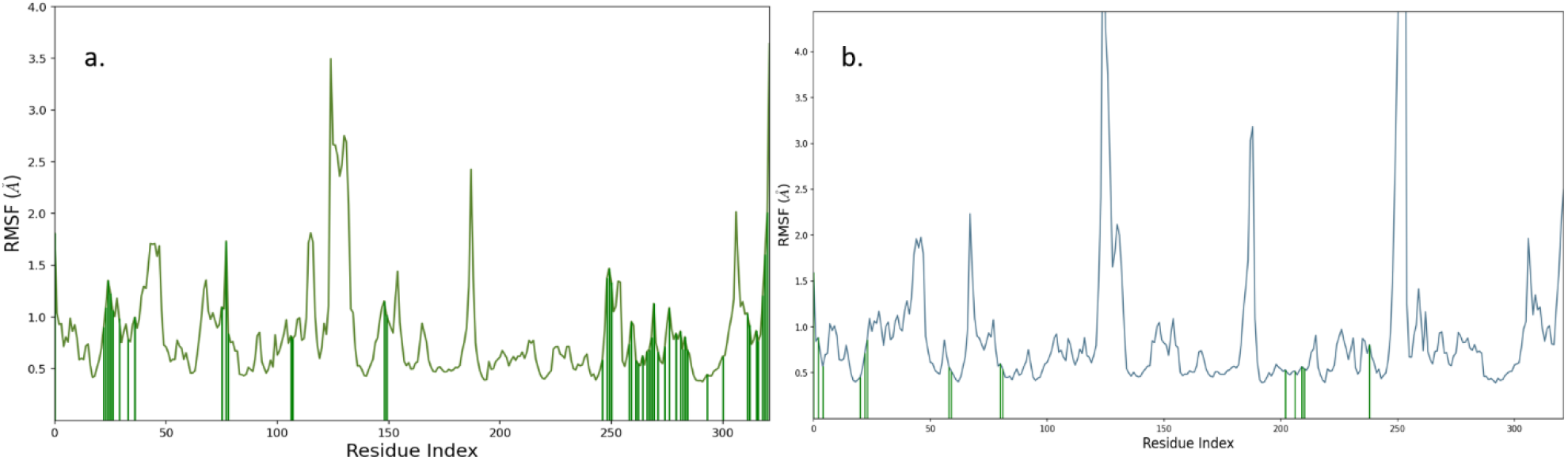
a. RMSF of Allosteric site residues during the 100ns simulation of 7S5H-Methylidene tetracyclo derivative. b. Allosteric site interaction formed during the 100ns simulation of 7S5H-Atrovastatin.

Fluctuation of residues were analyzed for both the complex in the allosteric site. The green colour line indicates the bond forming residues with the fragment of molecules. Figure a illustrates the fluctuation of residues by Methylidene tetracyclo derivative fragment binding. The amino acid residues from 31 to 40 made multiple interaction and the 246 to 340 residues holding the residues to make proper stable secondary structure. In the Atorvastatin, pyran fragment made few interactions with the allosteric site residues.

The overall work proved that Methylidene tetracyclo derivative is binding well in the active site and allosteric site compared to atorvastatin.

## 4. Conclusion

The results of interaction analysis and stability studies of this work concluded that the molecule Methylidene tetracyclo was better lead molecule to treat the atherosclerosis. Specifically, the tetrahydro pyran ring substituted with OH groups increase affinity to active and allosteric site amino acids. This leads the better possible drug molecule and the MD stability study confirms the unbreakable non-bonded interaction, which mimics the molecule is stable inside the protein to produce the complete malfunction. Fabrication of the rigid skeleton may increase the activity better than the Atorvastatin to treat the Atherosclerosis.

